# Breast Cancer Cells and Macrophages in a Paracrine-Juxtacrine Loop

**DOI:** 10.1101/2020.06.16.154294

**Authors:** Sevgi Onal, Merve Turker-Burhan, Gizem Bati-Ayaz, Hamdullah Yanik, Devrim Pesen-Okvur

**Affiliations:** Graduate Program in Biotechnology and Bioengineering, Gulbahce Kampusu, Urla, Izmir 35430, Turkey; Department of Molecular Biology and Genetics, Izmir Institute of Technology, Gulbahce Kampusu, Urla, Izmir 35430, Turkey

**Author notes:** MacDiarmid Institute for Advanced Materials and Nanotechnology, Electrical and Computer Engineering, University of Canterbury, Christchurch 8020, New Zealand (Sevgi Onal). Izmir International Biomedicine and Genome Institute, Dokuz Eylul University, Izmir 35330, Turkey (Merve Turker-Burhan). Deparment of Basic Oncology, Hacettepe University Cancer Institute, Ankara 06230, Turkey (Hamdullah Yanik). Corresponding author: Devrim Pesen-Okvur, Department of Molecular Biology and Genetics, Izmir Institute of Technology, Gulbahce Kampusu, Urla, Izmir 35430, Turkey.

## Abstract

Breast cancer cells (BCC) and macrophages are known to interact via epidermal growth factor (EGF) produced by macrophages and colony stimulating factor-1 (CSF-1) produced by BCC. Despite contradictory findings, this interaction is perceived as a paracrine loop. Further, the underlying mechanism of interaction remains unclear. Here, we investigated interactions of BCC with macrophages in 2D and 3D. BCC did not show chemotaxis to macrophages in custom designed 3D cell-on-a-chip devices, which was in agreement with ELISA results showing that macrophage-derived-EGF was not secreted into macrophage-conditioned-medium. Live cell imaging of BCC in the presence and absence of iressa showed that macrophages but not macrophage-derived-matrix modulated adhesion and motility of BCC in 2D. 3D co-culture experiments in collagen and matrigel showed that BCC changed their multicellular organization in the presence of macrophages. In custom designed 3D co-culture cell-on-a-chip devices, macrophages promoted and reduced migration of BCC in collagen and matrigel, respectively. Furthermore, adherent but not suspended BCC endocytosed EGFR when in contact with macrophages. Collectively, our data revealed that macrophages showed chemotaxis towards BCC whereas BCC required direct contact to interact with macrophage-derived-EGF. We propose that the interaction between cancer cells and macrophages is a paracrine-juxtacrine loop of CSF-1 and EGF, respectively.

## Introduction

Metastasis is the leading cause of death for cancer patients. As cancer cells metastasize, they interact with various extracellular molecules, growth factors and stromal cells such as macrophages and fibroblasts [1, 2].

Growth factors act as intercellular signaling molecules that promote various processes such as cell growth, adhesion and motility. Growth factors can be soluble, transmembrane or extracellular matrix bound proteins [3, 4]. Epidermal growth factor (EGF) is one of the seven ligands of EGF receptor (EGFR also known as ErbB1), and is the most studied member of the ErbB receptor family. While other EGFR ligands can bind to different members of the ErbB family, EGF binds only to EGFR [5-7]. In addition, EGFR expression correlates with poor prognosis in breast cancer [8, 9]. Mature EGF (6 kDa) is not detected in conditioned medium, suggesting that EGF is not secreted and direct contact may be required [10, 11]. It is also known that soluble EGF and conditioned medium of macrophages do not promote breast cancer cell invasion into collagen matrix and breast cancer cells do not invade into collagen if they are not co-cultured with macrophages[12]. Furthermore, it has been shown that EGFR can be activated with membrane bound ligands [13, 14]. Macrophage colony-stimulating factor (CSF-1) is known to regulate the proliferation, differentiation, survival and motility of macrophages [15]. Other ligands such as the EGF-like ligand Heregulin β1 (HRGβ1) binding to ErbB3 or ErbB4, and CXCL12 binding to CXCR4 abundant on invasive breast cancer cells relies on EGF and CSF-1 interaction to induce breast cancer metastasis *in vivo* [16].

Macrophages are stimulated towards the tumor micro-environment by growth factors and chemokines, for example, CSF-1. Macrophages have been shown to promote tumor growth; facilitate angiogenesis, lymphangiogenesis, stromal remodeling; change multicellular organization of cancer cells; induce invasion and metastasis [17-20]. Tumor-associated macrophages support the migration of cancer cells by the growth factors they express, for example, EGF. Thus, macrophages and their interactions with cancer cells are promising targets to work on for discovery of new therapeutic agents and approaches to manage cancer metastasis. To achieve this goal, the underlying mechanism of interactions between macrophages and cancer cells needs to be well-defined. While their interactions have been perceived as a paracrine loop of EGF and CSF-1 [21, 22], an in-depth understanding of the mechanistic basis of this interaction is lacking.

Most widely used *in vitro* cell culture systems neither reflect the organization and complexity of the *in vivo* microenvironment nor provide extensive spatial and temporal control. On the other hand, microfluidics based cell-on-a-chip devices can provide both 2D and 3D settings, position multiple cell types at specific locations, provide static and dynamic chemical and physical inputs and gradients, mimic physiologically relevant cell-to-cell and cell-to-matrix interactions and enable real time monitoring or visualization [23-27]. Therefore, cell-on-a-chip devices are now proving to be a necessary step which links *in vitro* studies, *in vivo* animal models and clinical trials.

Here, using a multidisciplinary approach including classical and state-of-the-art techniques such as live cell imaging and cell-on-a-chip devices, we showed that the interaction between BCC and macrophages is a paracrine – juxtacrine loop and direct contact is required for the activity of macrophage-derived-EGF on breast cancer cells.

## Results

### BCC did not show chemotaxis towards macrophages whereas macrophages showed chemotaxis towards BCC

To determine the mechanism of interaction between macrophages and BCC on the EGF – CSF-1 axis, in particular to determine how macrophage-derived-EGF acts on BCC, we first investigated chemotaxis in 3D cell culture. To assess invasion and migration capacity of breast cancer cells (BCC) and macrophages (MC), we first used Invasion-Chemotaxis chips (IC-chips) with three tandem channels (Fig. 1A). Here, constituents from adjacent channels had access to each other through gaps between regularly spaced posts that formed the borders between channels. Cell-free growth factor reduced matrigel was loaded into the middle channel. After matrix polymerization, either culture medium supplemented with 10% fetal bovine serum (FBS) or serum free medium was loaded into the chemoattractant (bottom) channel. Finally, BCC or MC suspended in serum free medium were loaded to the cell (top) channel. The chips were incubated upright to allow cells settle down at the medium-matrix interface and start invasion and migration at the same borderline. IC-chips were imaged at day 1 and day 3 using confocal microscopy (Fig. 1B). Image analysis showed that both BCC and MC showed prominent migration towards FBS but not serum free medium (p<0.05) (Fig. 1C).

**Figure 1.**
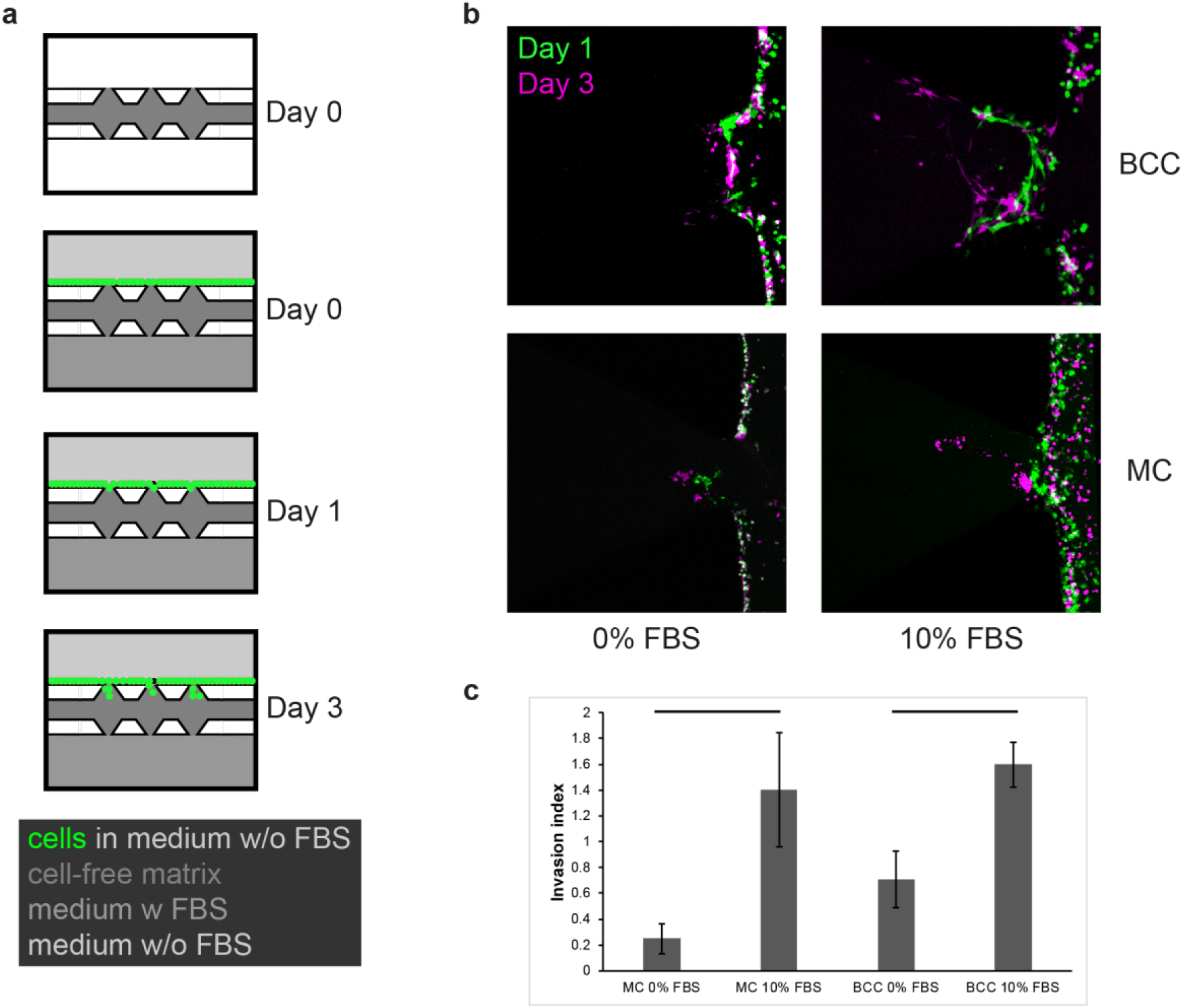
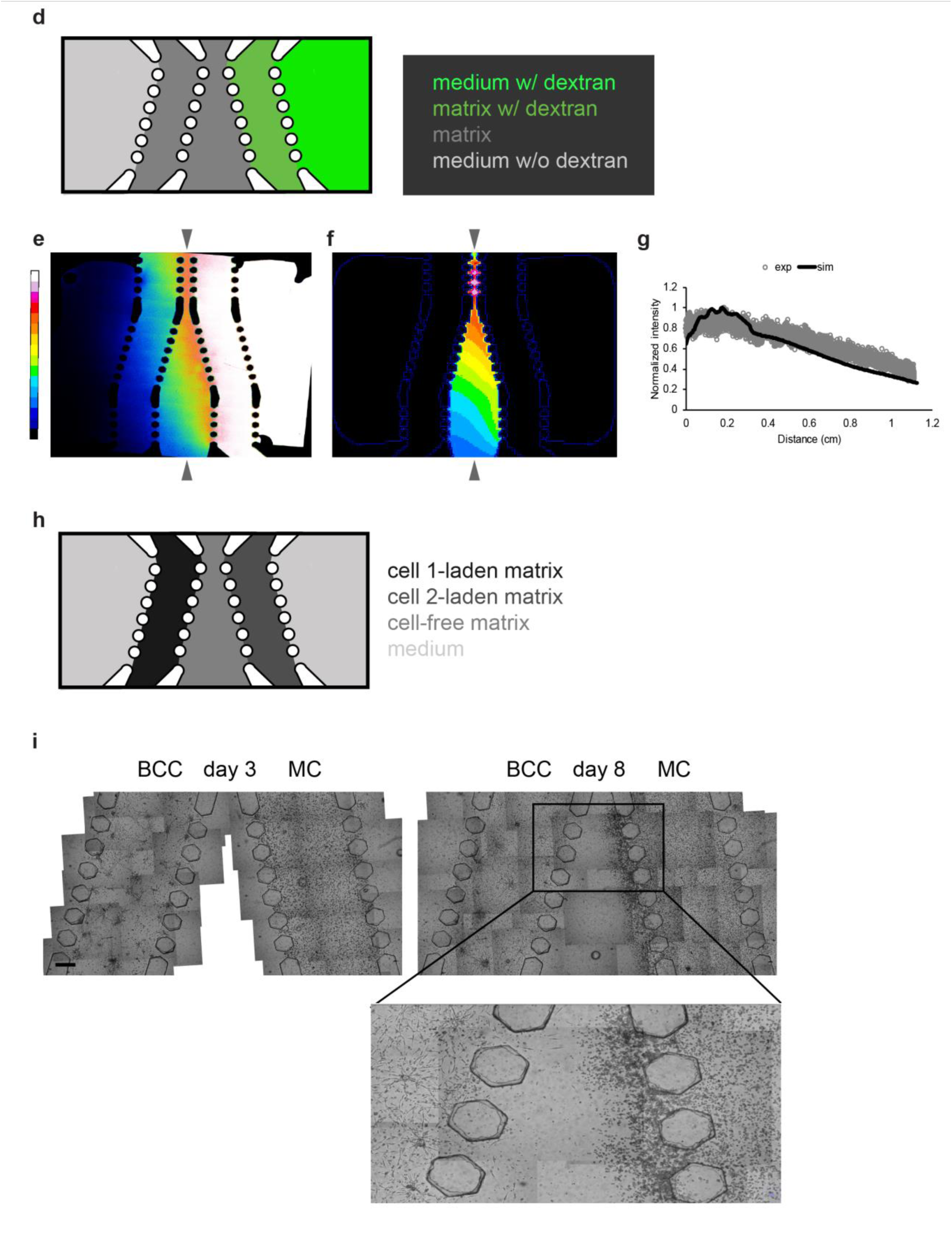
BCC cells did not show chemotaxis towards MC whereas MC showed chemotaxis towards BCC. (a) Cell-on-a-chip design (IC-chip) to test invasion and migration capacity of BCC and MC (not drawn to scale). Cell-free matrix was loaded into the middle channel. Either culture medium with FBS or serum-free medium was loaded into the bottom channel. Cells suspended in serum-free medium were loaded to the top channel. (b) Confocal images of BCC and MC at the medium-matrix interface and in the matrix across the serum-free medium (0% FBS) or culture medium with 10% FBS as chemoattractant on Day 1 and Day 3. (c) Prominent migration of MC and BCC towards FBS but not serum free medium. Horizontal bars show significant differences between the serum-free and FBS groups for each cell type (mean ± s.e.m. n = 3-13). (d) Cell-on-a-chip design (DDI-chip) to test the diffusion of the dextran molecule (not drawn to scale). Cell-free matrix was loaded into the middle channel. Dextran-laden matrix was loaded into the side channel adjacent to the middle channel. Dextran-free matrix was loaded into the other side channel. The reservoir neighbouring the dextran-laden matrix channel was filled with medium containing dextran. The other reservoir neighbouring the free matrix was filled with dextran-free medium. (e) Fluorescence image of the diffusion of 10 kDa fluorescent dextran in the DDI-chip at day one. (f) Simulation result of the diffusion of 10 kDa dextran molecule in the DDI-chip at day one, generated by VCell. (g) Gradient profiles of the dextran molecule along the distance marked by grey arrowheads in the experimental and simulation results. (h) Cell-on-a-chip design (DDI-chip) to test distant interactions (not drawn to scale). Cell-free matrix was loaded into the middle channel. Cell-laden matrices were loaded into channels on either side of the middle channel. The two reservoirs neighbouring the cell-laden channels were filled with cell culture medium. (i) Representative image for a DDI-chip loaded with BCC and MC (n = 6 cell-on-a-chip devices). (Scale bars, 500 µm.)

To evaluate chemotaxis between cells, we used a custom cell-on-a-chip device comprising a total of five tandem channels again connected to each other with regularly spaced posts. We loaded cell-free matrix into the middle channel (2 mm width) and then different cell-laden matrices into the channels at the left and right of the middle channel. The two outermost channels, were filled with serum free culture medium. Such a cell-on-a-chip design allowed assessment of the chemotactic responses between two cell types in a 3D cell culture setting. Here, macrophages showed low level of migration towards BCC which, on the other hand, did not migrate (Fig. S1A, B). To remove any limitations due to the absence of serum and long distances between cells, we used the Distance Dependent Interactions chips (DDI-chips) where the distance between the two cell types changed from 0.3 mm to 3 mm, and the cell culture medium in the reservoirs contained serum [28]. We first examined diffusion of 10 kDa fluorescent dextran in the DDI-chip experimentally. Fluorescent microscopy images acquired after one day showed that a gradient of the dextran molecule formed in the DDI-chip, as expected (Fig. 1D, E). We then examined diffusion of 10 kDa dextran molecule in the DDI-chip using VCell [29]. Simulation results for the duration of one day showed formation of a gradient of the dextran molecule, in agreement with the experimental results (Fig. 1F, G, Supplementary Movie S1). Control experiments where only BCC or only macrophages or BCC across normal mammary epithelial cells were cultured in the DDI-chip showed no significant migration (Fig. S1C, D, E). However, when BCC and macrophages were across from each other in the DDI-chip, macrophages showed prominent migration towards BCC which still did not migrate notably (Fig. 1H, I). These results showed that BCC did not show chemotaxis towards macrophages whereas macrophages did so.

To confirm that BCC provided a soluble signal whereas macrophages did not, we determined the EGF and CSF-1 content of macrophage- and BCC-conditioned medium (CM), Macrophage- and BCC-derived-extracellular matrix (ECM) and the cells themselves using ELISA. The majority of the protein and the growth factors were present in cells, as expected (Table 1). The ECMs from macrophages and BCC constituted about 37% and 19% of the total protein and they contained 7% and 12% of EGF and CSF-1, respectively. The conditioned medium of macrophages was 1% of the total protein content and it contained only 1% of the total EGF, showing that EGF was not secreted. Yet, the conditioned medium of BCC was almost 1% of the total protein content and contained 35% of the total CSF-1 showing that CSF-1 was secreted. Concentration of EGF in macrophage-CM was 0.0009 ng/ml whereas that of CSF-1 in BCC-CM was 0.544 ng/ml. We also measured EGF content of Matrigel (Corning) where [EGF]_avg in Matrigel_ is given as 0.7 ng/ml by the manufacturer within the range of 0.5 – 1.3 ng/ml. In agreement, we found [EGF]_Matrigel_ to be 0.978 ng/ml, while there is no CSF-1 in matrigel. Together, cell-on-a-chip and ELISA results indicated that macrophages could show chemotaxis to BCC-derived-CSF-1 whereas BCC did not show chemotaxis to macrophages, consistent with the lack of EGF in macrophage-conditioned-medium.

**Table 1.**
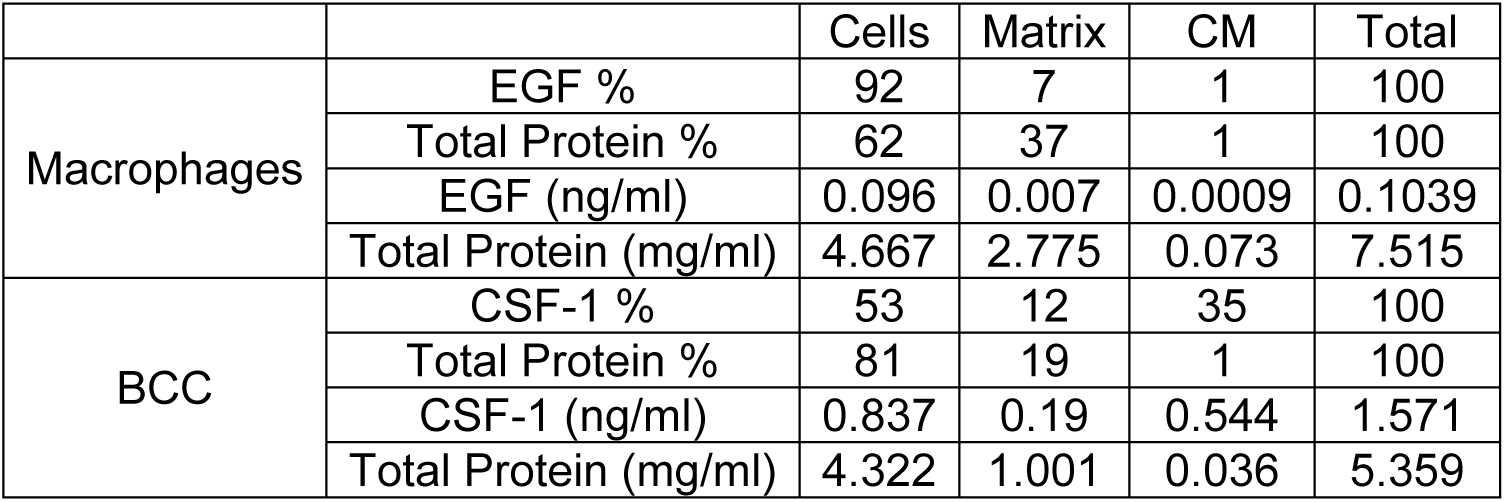
CSF-1 but not EGF was secreted. ELISA and total protein analysis for BCC, BCC-derived matrix, BCC-conditioned medium, MC, MC-derived matrix and MC-conditioned medium. Total % can exceed 100 due to rounding. The total protein and EGF as well as CSF-1 concentrations from which the percentages were derived are also given for the corresponding components of MC and BCC cultures.

### Macrophages but not macrophage-derived-matrix modulated adhesion and motility of BCC in an EGF-dependent manner

Since growth factors may bind ECM, we investigated adhesion and motility of BCC on macrophage-derived-ECM. BCC were imaged live as they were introduced onto glass coated with matrigel (mgel), glass coated with macrophage-derived-ECM (MCm), glass dispersedly coated with macrophages (MC) and bare glass surfaces. During the first fifty minutes, BCC on mgel surfaces attached and spread, increasing their cell area 4.79 fold (p<0.0001). Yet, BCC on the other surfaces did not spread significantly except on glass surface where there was a small (1.075 fold) increase in cell area (p<0.05). At fifty minutes, cell area on mgel surfaces was larger than those on all other surfaces (p<0.005) (Fig. 2A). Circularity of BCC decreased in time on mgel (p<0.001), but not on other surfaces. At fifty minutes, circularity of BCC on mgel surfaces was smaller than those on all other surfaces (p<0.001) (Fig. 2B). Aspect ratio of BCC did not change in time or between different surfaces (Fig. 2C). These results showed that presence of macrophages or macrophage-derived-ECM did not support initial cell attachment as well as matrigel.

**Figure 2.**
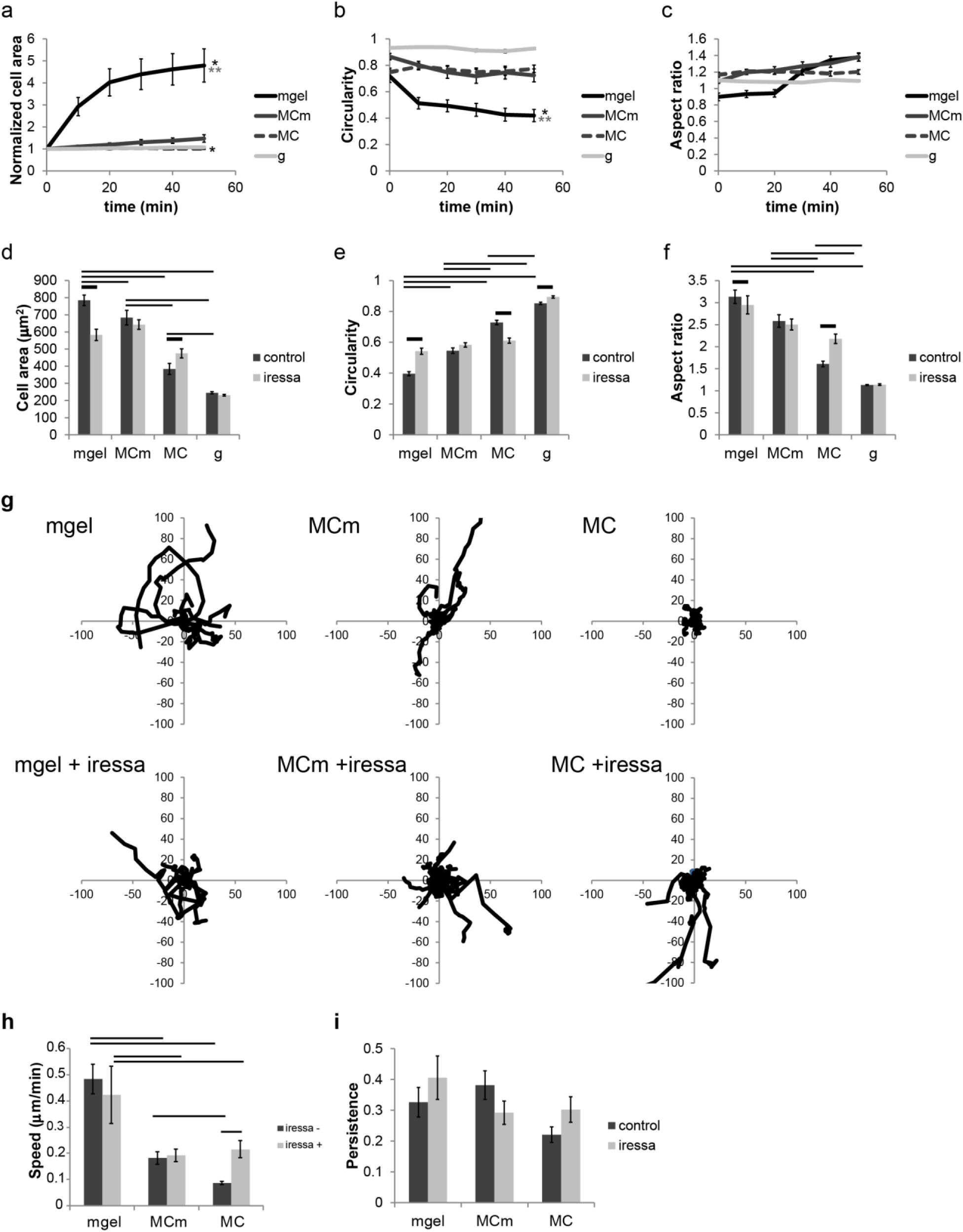
Macrophages but not macrophage-derived-matrix modulated adhesion and motility of BCC in an EGF-dependent manner. Quantification of (a) area, (b) circularity and (c) aspect ratio of cells during the first 50 minutes of adhesion (mean ± s.e.m. n = 18, 24, 23, 6 cells). Quantification of (d) area, (e) circularity and (f) aspect ratio of cells at 6 hours of adhesion in the presence and absence of iressa (mean ± s.e.m. n = 283, 145, 213, 97, 185, 255, 182, 130 cells). (g) Cell tracks of BCC motility on mgel, MCm, MC and glass surfaces in the absence and presence of iressa (IR) during 5 hours of live cell imaging (for n = 15-29 cells). Quantification of (h) average speed and (i) persistence of cells in the presence and absence of iressa (mean ± s.e.m. n = 20, 22, 29, 15, 24, 23 cells). Asterisks show significant differences between t = 0 and 50 minutes. Double asterisks show significant differences between matrigel and all other three surfaces. Horizontal bars show significant differences between control and iressa groups, all of which are not shown for clarity, but are available in Supplementary Excel File 1.

What is more, we analyzed cell morphology at the end of 5 hours on each of the above mentioned surfaces in the presence and absence of iressa (gefitinib), an EGFR inhibitor [30]. Areas of BCC decreased from mgel (784.5±30.9 μm^2^) to MCm (704.1±58.9 μm^2^) to MC (383.5±32.3 μm^2^) to glass (245.1±6.6 μm^2^) surfaces (p<0.036). Although the addition of iressa did not change the cell area of BCC on MCm and glass surfaces, it decreased and increased cell area on mgel (0.74 fold) and MC (1.24 fold) surfaces, respectively (p<0.0001) (Fig. 2D). Mgel is a rich surface as shown by effective adhesion of cells on it in the first fifty minutes unlike the other surfaces tested: The average cell area on mgel surfaces at fifty minutes was 1661.9±302.5 μm^2^, which was interestingly smaller than that at the end of 5 hours suggesting cells might be exploring less at later time points (784.5±30.9 μm^2^) (p<0.016). Yet, mgel surfaces allowed cell adhesion to mature and presence of iressa reduced cell area on mgel surfaces to (582.6±32.7 μm^2^) (p<0.0001). While the effect of iressa on cell area on MC surface may seem counterintuitive, the experimental set-up here is different in the sense that adhesion is examined in the first five hours of cells being introduced to a surface unlike many examples in the literature where adhesion and effect of iressa is examined in mature cultures. EGF is supposed to promote motility only when the pre-requirement of adhesion is satisfied. Here, BCC barely adhered on MC surfaces in the initial fifty minutes and had 2-fold smaller cell area compared to mgel surfaces at the end of 5 hours. Thus when the input for motility is quenched by iressa, cells could adhere better starting from suspended cells in medium. Circularity of BCC increased from mgel to MCm to MC to glass (p<0.0001). Presence of iressa increased the circularity of BCC on mgel and glass surfaces whereas it decreased that on MC (p<0.0001) surfaces (Fig. 2E). Aspect ratio of BCC was similar between mgel and MCm and decreased from MCm to MC to glass surfaces (p<0.0001). Presence of iressa decreased and increased aspect ratio of BCC on mgel and MC surfaces, respectively (p<0.016) (Fig. 2F). Cell area, circularity and aspect ratio changes were also consistent with each other as less adherent cells tend to be more circular and have a smaller aspect ratio. These results showed that the presence of macrophage-derived-ECM supported adhesion and spreading of BCC as well as matrigel and better than the presence of macrophages at the end of five hours. Presence of iressa affected adhesion on mgel and MC but not MCm surfaces suggesting that EGF was present in matrigel and on macrophages.

Furthermore, we examined BCC motility on mgel, MCm and MC surfaces in the presence and absence of iressa during the first 5 hours of being introduced onto the surfaces of interest (Fig. 2G - I). The experimental set-up here is different in the sense that motility is examined in the initial hours of being introduced to a surface unlike many examples in the literature where motility is examined in mature cultures. Average speed of BCC on mgel (0.48±0.06 μm/min) surfaces was larger than those on MCm (0.18±0.02 μm/min) and MC (0.09±0.01 μm/min) surfaces (p<0.00002). Iressa did not have an effect on average speed of BCC on mgel most likely because the rich composition of matrigel provided compensation. Iressa did not change average speed of BCC on MCm surfaces as well, considering cells on MCm surface already had low motility, their average speed did not change probably because at that stage the cells could not effectively utilize EGF signaling, because the pre-requirement for adhesion was not satisfied. Cells can be motile only after they have adhered well enough and thus there is a positive feedback from adhesion to motility. Thus MCm surfaces promoted cell adhesion but not motility. Yet, presence of iressa increased the average speed of BCC on MC surfaces 2.5 fold (p<0.00001), which was consistent with the increase in cell adhesion in the presence of iressa on MC surfaces. When iressa is present, the EGF induced motility signaling is quenched and the cells have a chance to adhere first. Consequently, with increased adhesion, the adhesion prerequisite for motility is satisfied and motility can increase. Lastly, persistence of BCC on all surfaces was similar (Fig. 2I). Finally, any EGF mediated effect on cell adhesion and motility was apparent on MC but not MCm surfaces. These results aligned with ELISA results showing majority of EGF was associated with macrophages.

### Macrophages promoted and reduced migration of BCC in collagen and matrigel, respectively

As cells can also interact with membrane-bound growth factors, it is possible that BCC interact with EGF which is macrophage-bound. In this case, direct contact with macrophages is likely to modulate phenotypes of BCC. Results for adhesion and motility of BCC on MC surfaces reported above supported such a juxtacrine mode of interaction. Here, we further investigated BCC and macrophages in 3D co-culture (Fig. 3 and Fig. 4). The multicellular organization of BCC changed in collagen and matrigel hydrogel drops in the presence of macrophages. In collagen, BCC appeared as round or elongated and along or elongated and perpendicular cells as well as clusters along the cell-laden hydrogel drop border. On day 5 of co-culture, presence of macrophages changed the percentile distribution of these structures (χ^2^ p<5.77303E-14). Percentage of along and clustered cells decreased and increased, respectively (Percent t-test <0.05). The number of round cells and clusters per hydrogel drop decreased (1.9-fold) and increased (24-fold), respectively (p<0.041). In matrigel, BCC alone organized into star-like multicellular complexes, branched structures or lines of cells. On day 5 of co-culture, presence of macrophages changed the percentile distribution of these structures (χ^2^ p<0.002). Percentage of branch and line structures decreased and increased, respectively (Percent t-test <0.05). The number of branched structures decreased 3-fold per hydrogel drop (p<0.029). Thus 3D co-culture results showed that BCC and macrophages did interact, resulting in changes in single and multi-cellular organization in 3D.

**Figure 3.**
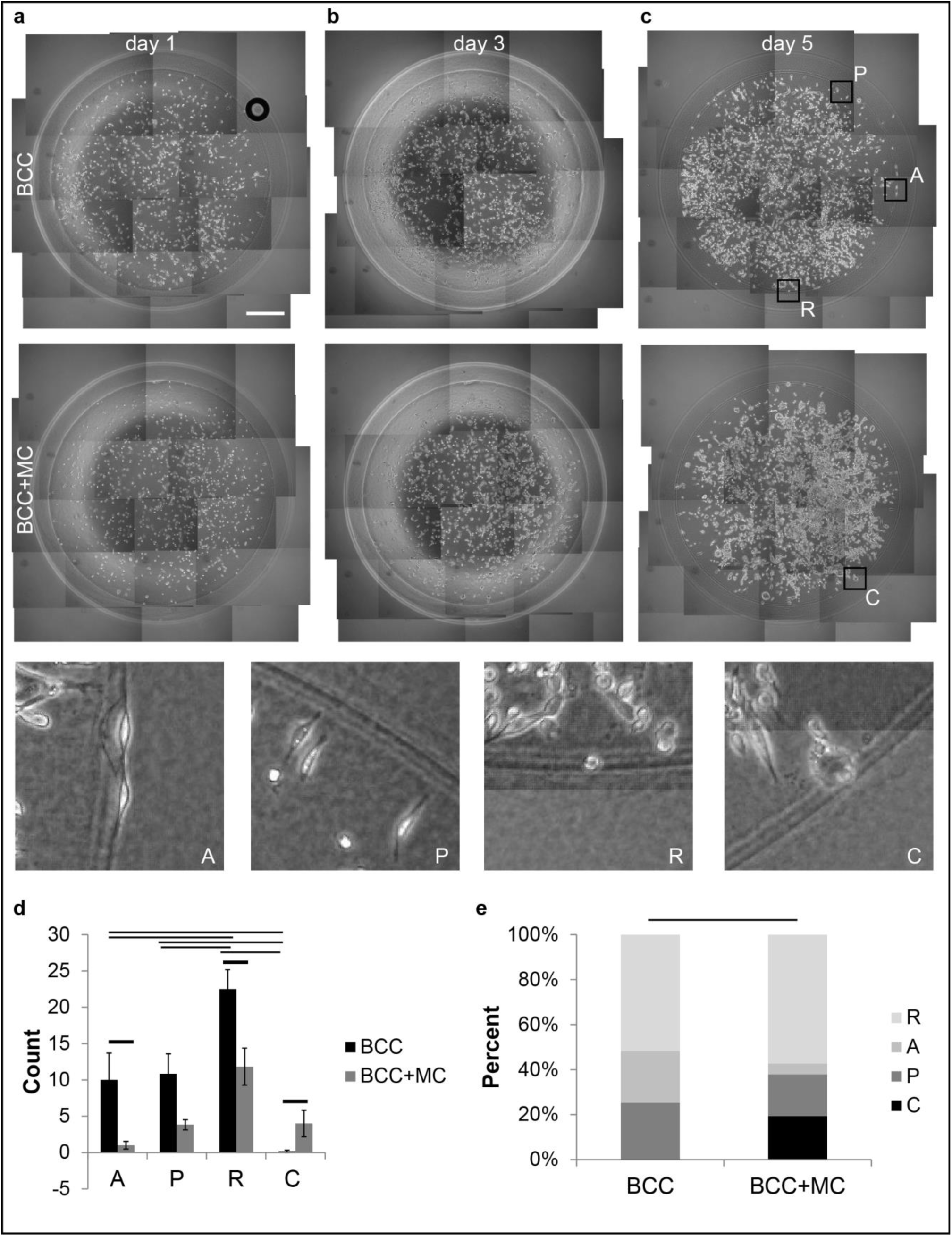
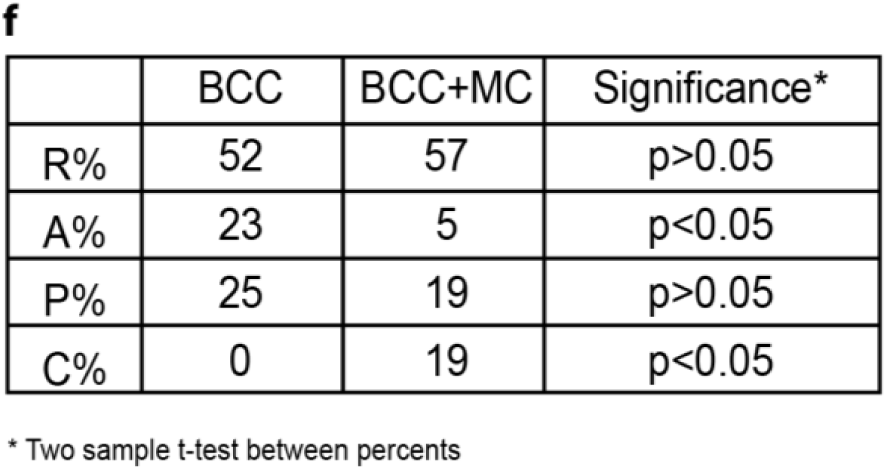
Co-culture of BCC with macrophages in collagen changed their multicellular organization. Presence of macrophages decreased the number of round cells (p<0.015) and increased the number of clusters per hydrogel drop (p<0.041), respectively and changed the percentile distribution of structures (χ^2^ test p<5.77E-14). The organization of BCC alone or with the presence of macrophages in collagen hydrogel drops on day 1 (a), day 3 (b) and day 5 (c). (Scale bars, 500 µm.) A: elongated and along, P: elongated and perpendicular, R: round, C: clusters along the cell-laden hydrogel drop border. (d) The number of the A, P, R, C structures on BCC alone and BCC co-culture with MCC on day 5 (mean ± s.e.m. n = 261, 124 structures). (e) The percentile distribution of the structures (χ^2^ test). Horizontal bars show significant differences. (f) Significances of the changes in the individual percentiles of R, A, P, C structures of BCC cultured in collagen alone or in the presence of macrophages (Two sample t-test between percents).

**Figure 4.**
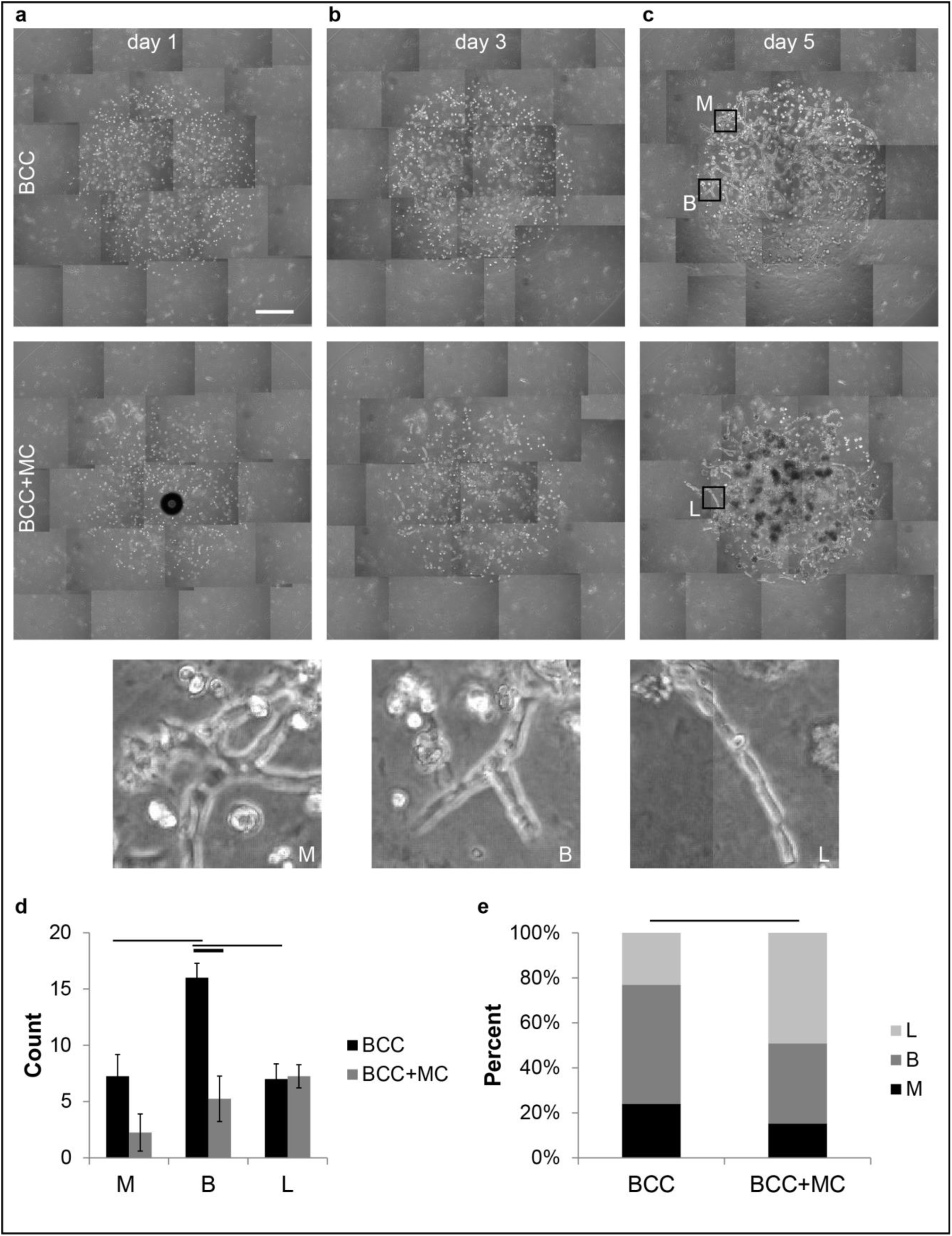
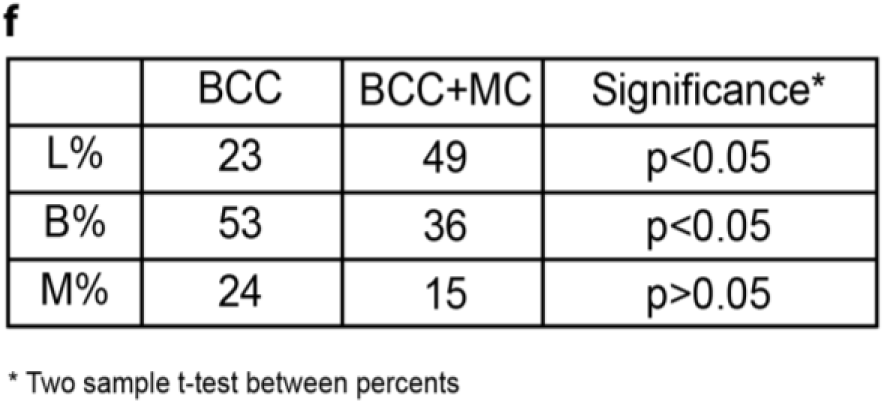
Co-culture of BCC with macrophages in matrigel changed their multicellular organization. Presence of macrophages decreased the number of branched structures of BCC per hydrogel drop 3-fold (p<0.029) and changed the percentile distribution of structures (χ^2^ test p<0.002). The multicellular organization of BCC in matrigel hydrogel drops alone or with the presence of macrophages on day1 (a), day 3 (b) and day5 (c). (Scale bars, 500 µm.) M: star-like multicellular complexes, B: branched structures, L: lines of cells. (d) The number of the M, B, L structures for BCC alone and BCC co-culture with MCC on day 5 (mean ± s.e.m. n = 121, 59 structures. (e) The percentile distribution of the structures (χ^2^ test). Horizontal bars show significant differences. (f) Significances of the changes in the individual percentiles of L, B, M structures of BCC cultured in matrigel alone or in the presence of macrophages (Two sample t-test between percents).

To determine cell migration in 3D in a more controlled manner, we used a custom 3D co-culture cell-on-a-chip device, where we seeded BCC or macrophages alone or in combination in collagen or matrigel into a channel sided by channels containing cell-free hydrogels (Fig. 5). 5% FBS supplemented RPMI medium was used in the medium reservoirs adjacent to the cell-free hydrogels to retain the 3D co-culture on-chip over 5 days. In collagen, both mono- and co-cultured cells showed increased migration from day 1 to day 3 to day 5 (p<0.05). What is more, BCC alone showed less migration than macrophages alone and presence of macrophages increased the migration distance 2.8 fold on day 5 (p<1.54E-06). In matrigel, both mono- and co-cultured cells showed increased migration from day 1 to day 5 (p<0.05). Migration of macrophages was significantly lower in matrigel than that in collagen (p<0.005). Furthermore, BCC alone showed more migration than macrophages alone and presence of macrophages reduced the migration distance 2 fold on days 1, 3 and 5 (p<0.028). Thus macrophages promoted and reduced migration of BCC in collagen and matrigel, respectively.

**Figure 5.**
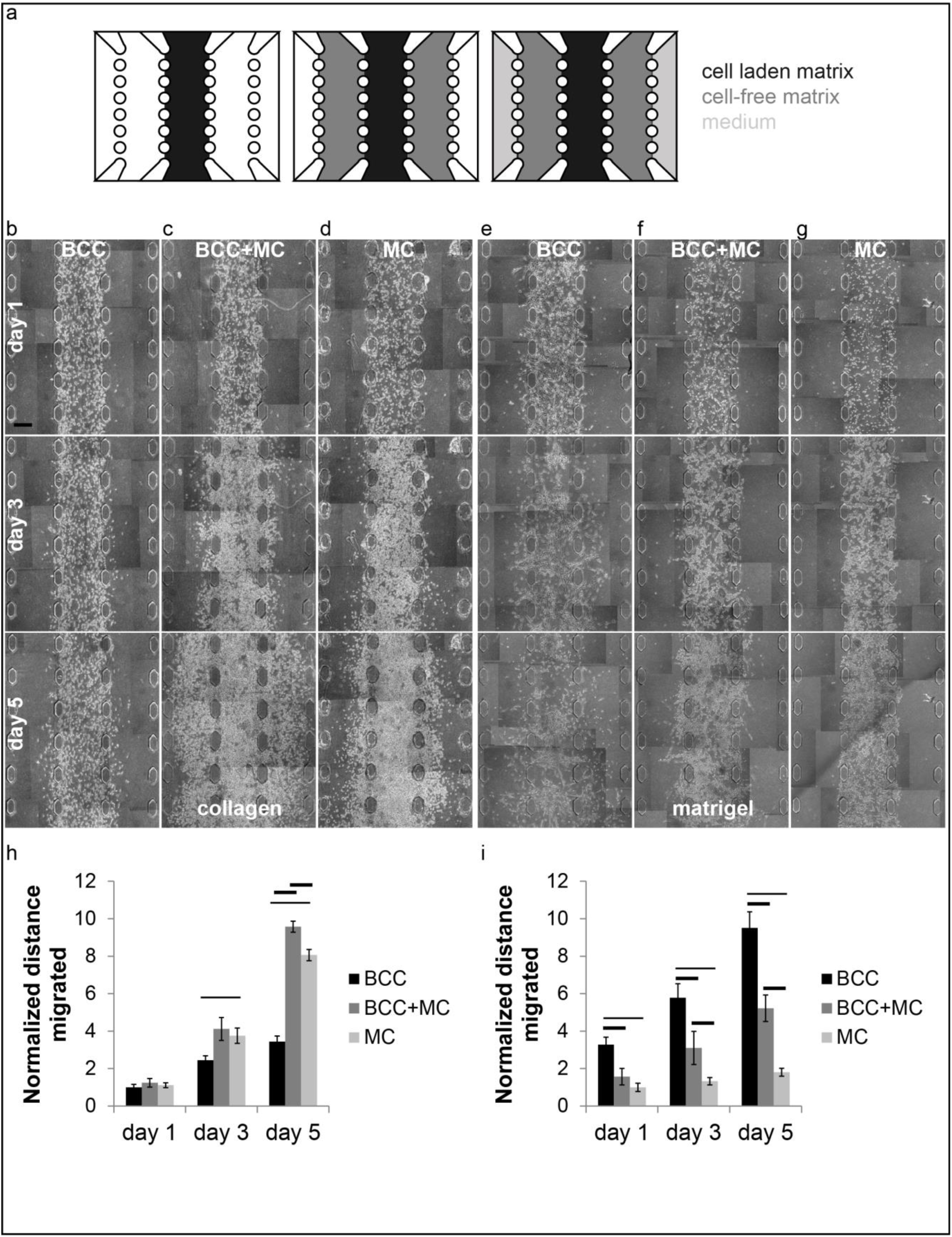
Macrophages promoted and reduced migration of BCC in collagen and matrigel, respectively. (a) Cell-on-a-chip design to test migration of alone and co-cultures of BCC and macrophages (not drawn to scale). Cell-laden matrices were loaded into the middle channel. Cell-free matrices were loaded into the adjacent channels on both sides of the middle channel. The reservoir channels neighbouring the cell-free hydrogel channels were filled with cell culture medium. (b) – (d) BCC alone, BCC and macrophages or macrophages alone in collagen were loaded into the middle channel of a cell-on-a-chip device. (e) – (g) BCC alone, BCC and macrophages or macrophages alone in matrigel were loaded into the middle channel of a cell-on-a-chip. Cell-free channels were loaded with the corresponding matrices. Quantification of distances migrated by cells in collagen (h) and matrigel (i) matrices (mean ± s.e.m. n = 16, 8 ROIs). Horizontal bars show significant differences between groups, all of which are not shown for clarity, but are available in Supplementary Excel File 1. (Scale bars, 250 µm.)

### Adherent but not suspended BCC endocytosed EGFR when in contact with macrophages

To confirm that juxtacrine signaling is the mechanism of interaction between macrophage-derived-EGF and BCC, we examined endocytosis of EGFR in BCC in suspension and adherent cell culture (Fig. 6). When starved BCC were treated with BSA, EGF or macrophages in suspension, the fraction of membrane EGFR was the highest for BCC treated with macrophages than with BSA than with EGF (p<0.0015) (Fig. 6A, B). EGFR was expected to be internalized in the presence of macrophage-derived-EGF. Yet interactions of BCC with macrophages did not promote receptor internalization, which was probably because BCC in suspension did not have enough traction to disengage the macrophage-bound-EGF, in agreement with previous work [31]. In adherent culture on the other hand, BCC cells transfected with EGFR-GFP starved and treated with macrophages endocytosed EGFR (69% of cells) more and less than those treated with BSA (11% of cells) and EGF (92% of cells), respectively (χ^2^ p<0.035) (Fig. 6C, D and Supplementary Movie S2, S3, S4).

**Figure 6.**
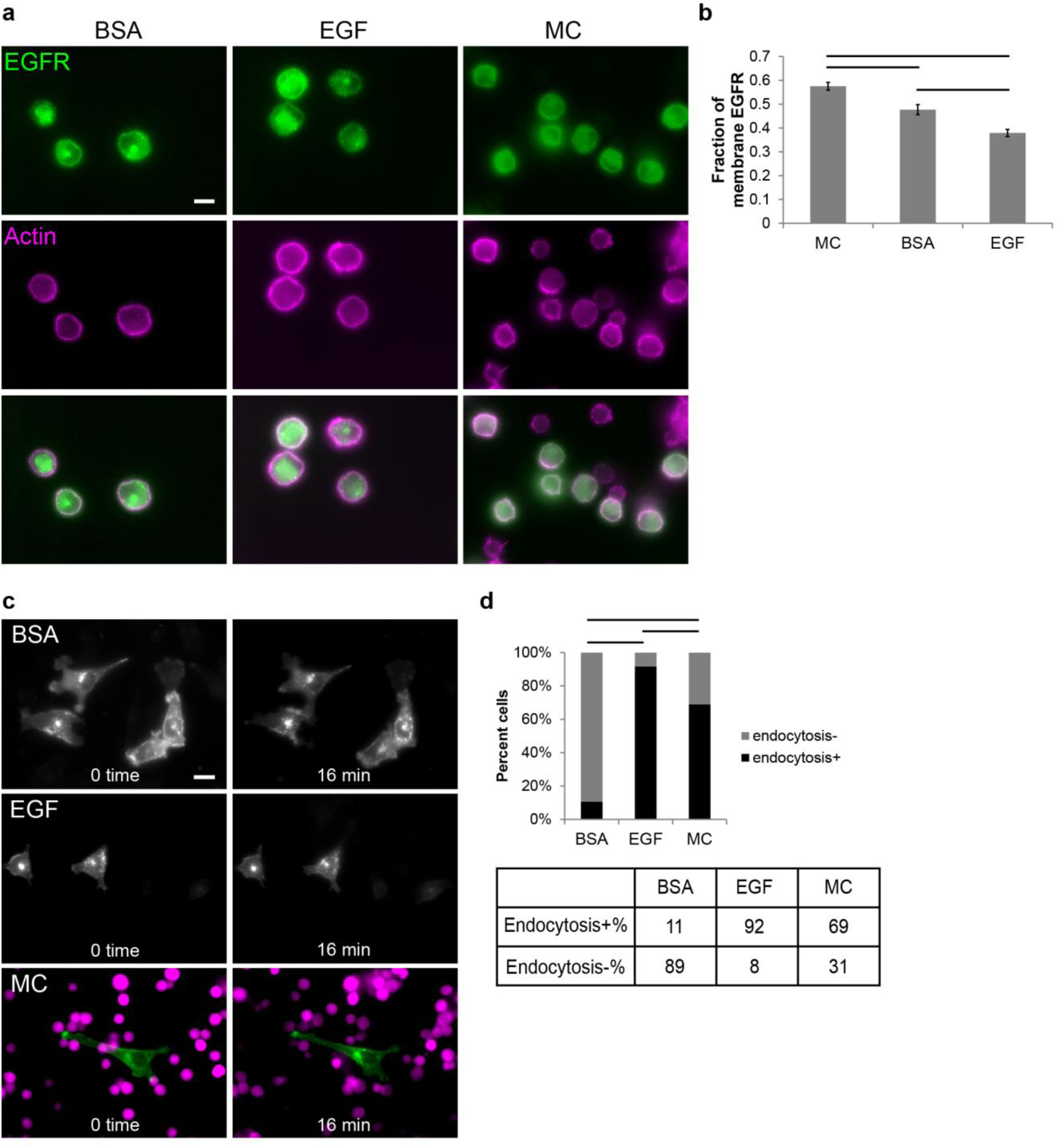
Adherent but not suspended BCC endocytosed EGFR when in contact with macrophages. (a) Starved and suspended BCC were treated with BSA, EGF or macrophages for 15 minutes in suspension, fixed and stained. Representative immunostaining images for EGFR and actin localization. (Scale bars, 10 µm.) (b) The fraction of membrane EGFR derived from immunofluorescence signal (mean ± s.e.m. n = 35, 45, 27 cells). (c) Representative images for 0th and 16th minute of live imaging of EGFR endocytosis in adherent BCC transfected with EGFR-GFP, starved and treated with EGF or macrophages. (Scale bars, 10 µm.) (d) The percentage of the BCC cells showing EGFR endocytosis when treated with BSA, EGF or macrophages (χ^2^ test for n = 66, 24, 42 cells). Horizontal bars show significant differences.

## Discussion

Although breast cancer cells (BCC) and macrophages are accepted to interact in a paracrine loop of epidermal growth factor (EGF) and colony stimulating factor-1, direct evidence to support this perception is lacking and the underlying mechanism of interaction remains unclear. We investigated the interaction between BCC and macrophages using a multidisciplinary approach. Our results support the hypothesis that a juxtacrine interaction is required for the activity of macrophage-derived-EGF on breast cancer cells, and thus the interaction between cancer cells and macrophages is a paracrine-juxtacrine loop of CSF-1 and EGF, respectively (Fig. 7).

**Figure 7.**
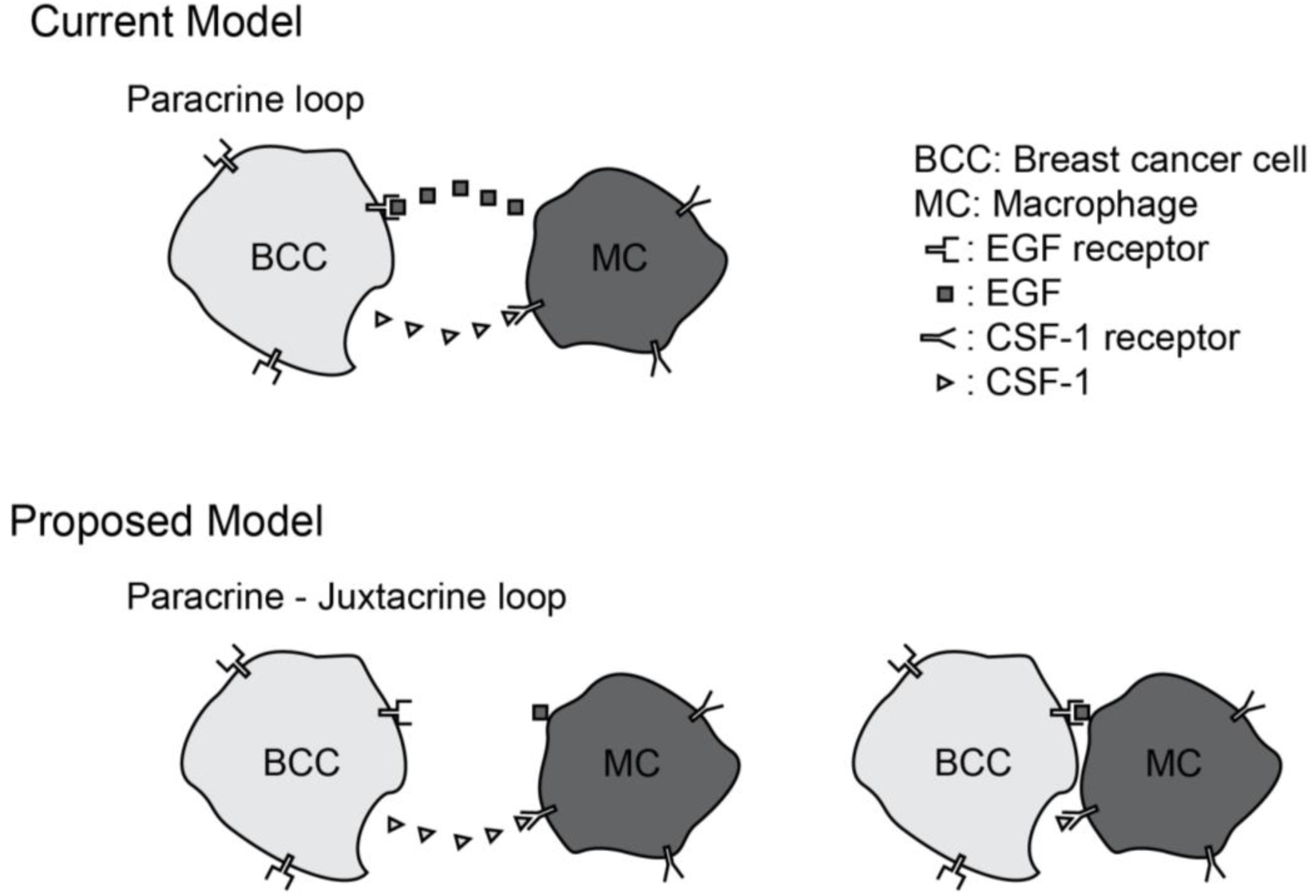
Current and proposed models for interaction of BCC with macrophages. In the current model (top), BCC show chemotaxis towards macrophage-derived-EGF and macrophages show chemotaxis towards BCC-derived-CSF-1. In the proposed model (bottom), macrophage-derived-EGF is associated with macrophages and direct contact is required for interaction of macrophage-derived-EGF and EGFR on BCC. Macrophages show chemotaxis towards BCC-derived-CSF-1, which is secreted.

Growth factors can act either in soluble or ECM-bound or cell-bound [7]. It should be noted that our experiments did not involve any exogenous EGF, the only source of EGF were the macrophages. Thus, we were able to examine cell-to-cell communication in a physiologically relevant microenvironment that mimicked the *in vivo* conditions. Future work could benefit from molecular knockdown of EGF, which is dispensable in the context of current work. Our first results showed that CSF-1 was secreted and thus a chemotactic response by macrophages towards BCC was possible and observed whereas EGF was not detected in the conditioned medium of macrophages and a chemotactic response by BCC to macrophage-derived-EGF was not observed. Secondly, we examined whether macrophage-derived-EGF could act as an ECM-bound growth factor. Here, we used mgel surfaces as positive controls. An important difference between mgel and MC surfaces was that unlike the latter, the former presented a rich ECM composition. Iressa decreased adhesion on mgel surfaces as expected since matrigel is a rich mixture of ECM proteins and growth factors. Presence of EGF can promote adhesion via crosstalk between integrins and growth factor receptors and presence of iressa can remove the pro-adhesion input from EGFR [32-35]. EGF is also known to promote motility. Macrophages appeared to inhibit cell adhesion and presence of iressa removed the pro-motility input from EGFR. This result was in agreement with the previous studies which found that EGF can promote rounding of adherent cells [36], inhibit adhesion [37] and promote a motile phenotype [38].

Adhesion of MDA-MB-231 cells, used here as a model for BCC, on collagen IV has been shown to increase in the presence of EGF and this increase can be reverted by EGFR inhibitors [39]. However, we cannot directly compare our results with those reported in that study because in our experimental system, soluble EGF is not present. Our results collectively indicated that macrophage-derived-EGF was cell-bound. On the other hand, in that study EGF has been shown to inhibit adhesion for cells with high EGFR expression. Thus it appears that the form of EGF – soluble or immobilized – and the number of EGFR per cell can modulate the effect of EGF on cell adhesion.

Iressa dependent differences on adhesion and motility were observed on macrophages but not on macrophage-derived-ECM, directing us to the investigation of cell-to-cell contact based interactions. In matrigel hydrogel drops, in the presence of macrophages, the number and percentage of branched structures decreased and the percentage of line structures increased suggesting that macrophages could induce a more dispersed organization of BCC. On the other hand, changes in the single and multi-cellular organization in collagen suggested that BCC and macrophages could cluster in a poor microenvironment such as collagen.

In 3D co-culture cell-on-a-chip devices, macrophages promoted and reduced migration of BCC in collagen and matrigel, respectively. However, it is possible that the effect of CSF-1 on macrophages is proliferation rather than migration. In collagen, BCC alone did not migrate as well due to the poor composition of collagen; whereas interactions with macrophages, which acted as rich sources of EGF, promoted cell migration, as expected. Our 3D migration results for cells in collagen in custom cell-on-a-chip devices are also in agreement with previous studies where dissemination of tumor cells is induced by contact with macrophages [12, 40]. Direct contact with macrophages is also known to induce other changes in cancer cells, such as formation of more invadopodia, which EGF is known to enhance [41]. On the other hand, in 3D co-culture cell-on-a-chip devices comprising matrigel, BCC alone could migrate well due to the rich composition of matrigel which can activate both integrins and growth factor receptors; yet as BCC encountered macrophages which acted as concentrated point sources of EGF, they migrated less. This was most likely because local amount of EGF, that was the sum of EGF present in matrigel plus macrophage-derived-EGF, became too high and inhibited migration of BCC, satisfyingly consistent with biphasic EGF dependence of EGFR auto-phosphorylation [42] and results of *in vivo* invasion assays performed with microneedles stably inserted into xenograft tumors in mice [43].

Our results on endocytosis of EGFR in suspension BCC when stimulated with macrophages are consistent with those of a study where cells were stimulated with surface immobilized EGF which has been suggested to be useful for studying juxtacrine signaling [44]. Furthermore, our results on endocytosis of EGFR in adherent BCC when stimulated with macrophages align with those of a study where cells were stimulated with EGF-beads [45]. These results are also in agreement with our ELISA results where EGF was detected with macrophages but not macrophage derived matrix or macrophage conditioned medium.

RAW264.7 is a commonly used cell line for convenience. In addition, macrophages are known to polarize into M1 and M2 phenotypes [46-48]. Using primary cells and examination of macrophage polarization under 3D co-culture conditions is beyond the scope of current work, yet, would be a focus of future work.

EGF – CSF-1 based interactions between cancer cells and macrophages have long been perceived as a paracrine loop. Using a multidisciplinary approach, our results revealed that cell-to-cell contact was required for the activity of macrophage-derived-EGF on BCC. To the best of our knowledge, this is the first study providing exhaustive evidence and showing that the mechanism of interaction between macrophage-derived-EGF and BCC is juxtacrine signaling. The paradigm shift we provide is likely to promote a better understanding of cell-to-cell communication in both health and disease states, and well-designed cellular microenvironments to control and assay cell-to-cell interactions in tissue engineering applications and finally better therapeutic and diagnostic approaches in the future. While our study reports novel results on the interactions of cancer cells and macrophages, the state-of-the-art cell-on-a-chip and 3D cell culture platforms developed here allow to use any cell, hydrogel and medium type of interest to study different cell-to-cell, cell-to-molecule, and cell-to-matrix interactions.

## Materials and Methods

### Cell culture

MDA-MB-231 (BCC), RAW264.7 macrophages and MCF10A were acquired from ATCC (LGC Standards GmbH, Germany). BCC and macrophages were grown in tissue culture treated petri dishes in DMEM supplemented with 10% FBS, 1X penicillin-streptomycin, 1X L-glutamine and in non-treated petri dishes in RPMI supplemented with 5% FBS, 1X penicillin-streptomycin, 1X L-glutamine, respectively, at 37°C, 5% CO_2_. BCC and macrophages were trypsinized and mechanically collected for sub-culturing, respectively. MCF10A were cultured as previously described [49].

### Cell-on-a-chip experiments

Fabrication of the cell-on-a-chip devices was performed as previously described [28] except that IC-chips (invasion-chemotaxis chips) were provided by Initio Biomedical (Turkey). In IC-chips, cell-free growth factor reduced matrigel (354230, Corning) was loaded into the middle channel. After matrix polymerization at 37°C and 5% CO2, either culture medium supplemented with 10% fetal bovine serum (FBS) or serum free medium, and either DsRed-labelled BCC or CellTracker Green stained MC suspended in serum free medium (1×10^6^ cells/ml) were loaded to the corresponding channels. The chips were incubated in a perpendicular orientation where the cells could flow downward onto the medium-matrix interface. IC-chips were imaged at day 1 and day 3 using a Leica SP8 confocal microscope. In other cell-on-a-chip devices, cell laden (6.5×10^6^ cells/ml) and cell-free matrigel (354234, Corning) or collagen gels (354249, Corning) were loaded to the corresponding channels and polymerized at 37°C and 5% CO2 for 30 min. Then culture media were loaded into the medium reservoirs. The samples were kept at 37°C and 5% CO_2_ for 7-14 days. Partially overlapping raster-scan phase-contrast images of fields of interest in cell-on-a-chip devices were acquired on at least days 1, 3 and 5 using an Olympus CX41 microscope or a Euromex OX.3120 microscope equipped with a Dino-Lite Eyepiece Camera and imaging software (DinoCapture 2.0). Images were stitched using Photoshop (Adobe).

For quantification of migration of co-cultured cells in cell-on-a-chip devices, each region between two PDMS posts was defined as an ROI and the maximum distance migrated in each ROI was measured using ImageJ/Fiji [50].

### Protein Quantification and ELISA

Macrophage-derived-matrix was prepared as described below. At least three biological and three technical repeats were carried out and representative results were reported.

1.158×10^6^ RAW 264.7 cells were seeded to get confluent macrophage cell matrix (MCm). The cells were cultured for 7 days before sample collection. 50% of old medium was replaced with fresh culture medium in every two days. At 5^th^ day, the old medium was replaced with fresh serum free culture medium and cells were cultured 2 days to produce macrophage conditioned medium. At 7^th^ day, the conditioned media was collected and filtered (0.2 µm PES) into a tube. 2M urea was used to remove cells. Urea supernatant including the detached cells was centrifuged at 400 rcf for 5 min. The cell pellet was suspended in 1X Diluent B of ELISA Kit (Abcam). After removing cells, the matrix remained in the petri dish was rinsed with 1X PBS four times. At last, the matrix was scraped and collected with 1X Diluent B and filtered (0.2 µm PES) into a tube. All of the samples were stored at −80°C till measuring their EGF content with EGF Mouse ELISA. MDA-MB-231 cells were processed similarly. All the samples were processed for Bradford (39222.02, Serva), EGF Mouse ELISA (ab100679, Abcam) and CSF Human ELISA (ab100590, Abcam) assays according to the manufacturers’ instructions.

### Live cell imaging

BCC were starved in serum free Leibowitz’s medium supplemented with BSA, collected using cell dissociation buffer (Biological Industries, Israel) and re-suspended in starvation medium and added on glass, matrigel, macrophage-derived-matrix or macrophages. Imaging was started immediately using an Olympus IX70 microscope equipped with a heating plate set to 37°C. Phase-contrast images were captured with a Euromex camera with the ImageFocus Software every 30 seconds.

For mgel surfaces, 100 μg/ml matrigel was used for coating glass coverslips. For MCm surfaces, macrophage derived matrix was prepared by seeding 48000 RAW 264.7 cells per 15mmx15mm area of a glass coverslip and culturing cells for 7 days prior to the live cell imaging experiment. Macrophages were removed using 2M urea. For MC surfaces, 6000 cells were seeded, cultured for 7 days and used after rinsing with serum-free medium.

For live cell experiments on MC surfaces, BCC and macrophages were stained with CellTracker Green CMFDA or Blue CMAC (Molecular Probes), respectively, according to the manufacturer’s instructions. Fluorescence images were captured for the first and last time points.

BCC were treated with 2 μM Iressa (‘Gefitinib’ sc-202166, Santa Cruz Biotechnology) for 16 hours prior to using the cells in live cell imaging experiments. Medium with Iressa was replenished just before live cell imaging.

Cell area, circularity and aspect ratio of the cells were measured from manually tracked cell boundaries using ImageJ. For motility, cell nuclei were manually tracked over time. Speed was calculated as the ratio of the net distance travelled to time for each time interval of 15 minutes. Persistence was calculated as the ratio of the net distance to the total distance.

### 3D Co-culture hydrogel experiments

2×10^6^ cells/ml of BCC and macrophages were seeded alone or together in 1:1 matrigel or 2 mg/ml collagen hydrogel drops of 2 µl in multi-well plates which were placed upside down during hydrogel polymerization. Another 15 µl of the corresponding cell-free hydrogel was then polymerized on the cell-laden hydrogels. Next, macrophage culture medium was added to the wells, and cells were cultured at 37°C and 5% CO_2_. Image acquisition was performed as for cell-on-a-chip experiments.

The outermost 328 μm (250 pixels) ring of the cell-laden matrigel drops was examined. A line structure was defined to contain at least 2 cells and be more than 100 pixels in length. A branch was defined to contain at least 3 cells and to have a ‘Y’ or ‘T’ shape. A multicellular complex was defined to contain at least 4 cells which had connections with each other.

The boundary at the cell-laden and cell-free collagen was examined. An along cell was defined to be aligned along the boundary. A perpendicular cell was defined to be perpendicular to the boundary. Round and clustered cells at the boundary were also counted.

Assignments of different structures were performed by two or three independent observers and cross-checked.

### Endocytosis in suspended cells

BCC were starved and incubated in a cell dissociation buffer (Biological Industries, Israel) for collection. BCC were then treated with 3.5 nM EGF or macrophages in suspension for 10 minutes. Samples were then fixed with 4% paraformaldehyde and processed for immunostaining with EGFR (D38B1) XP rabbit mAb (4267, Cell Signaling Technology, 1:100), anti-rabbit secondary antibody Alexa Fluor 555 Conjugate (4413, Cell Signaling Technology, 1:200) and Alexa Fluor 488 Phalloidin (8878, Cell Signaling Technology, 1:200). Fluorescence images were captured with an Olympus IX83 microscope equipped with a DP73 camera and CellSens software. Fluorescence signal of EGFR localized to the membrane divided by the total cellular signal was measured using ImageJ.

### Endocytosis in adherent cells

BCC were transiently transfected with EGFR-GFP, a gift from Alexander Sorkin (Addgene plasmid # 32751). BCC were starved and treated with 3.5 nM EGF or suspended macrophages labelled with Blue CMAC (Molecular Probes). Images were acquired with a Zeiss Observer microscope equipped with an incubation chamber set to 37°C, an MRm camera and Zen software. BCC showing inward movement of EGFR-GFP from the cell membrane to the cytosol were counted as endocytosis positive.

### Image analysis

Photoshop (Adobe) and ImageJ (NIH) were used for image processing and analysis.

### Diffusion in DDI-chip

Matrigel was diluted 1:1 with medium supplemented with 10 kDa fluorescent dextran (final concentration 5 µM) and loaded into the right matrix channel. Fluorescent dextran (final concentration 10 µM) was loaded into the right medium channel. Matrigel diluted 1:1 with medium was loaded into the middle and left channels. Medium was loaded into the left medium channel. Fluorescence images were acquired after one day.

### Simulation

VCELL [29] was used for simulation of diffusion of fluorescent 10 kDa dextran in the DDI-chip using the parameters in the diffusion in DDI-chip experiment. The model is available on request.

## Statistical Analysis and Data Presentation

Mann-Whitney two-tailed test (MATLAB), χ^2^ test (Microsoft Excel) and two sample t-test between percents (StatPac) were used to determine significant differences in mean and percentage values, respectively. Statistical significance was taken as p < 0.05. Data were represented as means ± s.e.m. All statistical test results are available as Supplementary Excel File 1. All data used for statistical analysis is available as Supplementary Excel File 2.

## Authors’ Contributions

Sevgi Onal: Investigation, Formal analysis, Validation, Visualization, Writing - Original Draft, Writing - review & editing

Merve Turker: Investigation, Formal analysis, Writing - Original Draft Gizem Bati: Investigation, Formal analysis, Writing - review & editing Hamdullah Yanik: Investigation

Devrim Pesen-Okvur: Conceptualization, Methodology, Investigation, Validation, Formal analysis, Visualization, Writing - Original Draft, Writing - review & editing; Supervision, Funding acquisition

## Acknowledgements

This work was supported by FP7 Marie Curie Grant number PIRG08-GA-2010-276976 and IYTE Scientific Research Project Grant number 2011IYTE25 (to Devrim Pesen-Okvur). Cell-on-a-chips were fabricated at the IYTE Applied Quantum Research Center, supported by DPT (State Planning Organization) Grant 2009K120860.

We thank E. Koc for assisting with English language editing, O. Yalcin-Ozuysal for critical reading of the manuscript, A. Arslanoglu, E. Ozcivici and Biotechnology and Bioengineering Application and Research Center for access to fluorescence microscopes, Z. Ulger and S. Yucel for help with manual cell tracking.

## Additional Information

Supplementary information accompanies this paper. Data available on request from the authors.

## Competing financial interests

Devrim Pesen-Okvur and Sevgi Onal were co-founders of INITIO Biomedical Engineering Consulting Ind. Tra. Ltd. Co., Izmir, Turkey.

---

**Figure S1.**
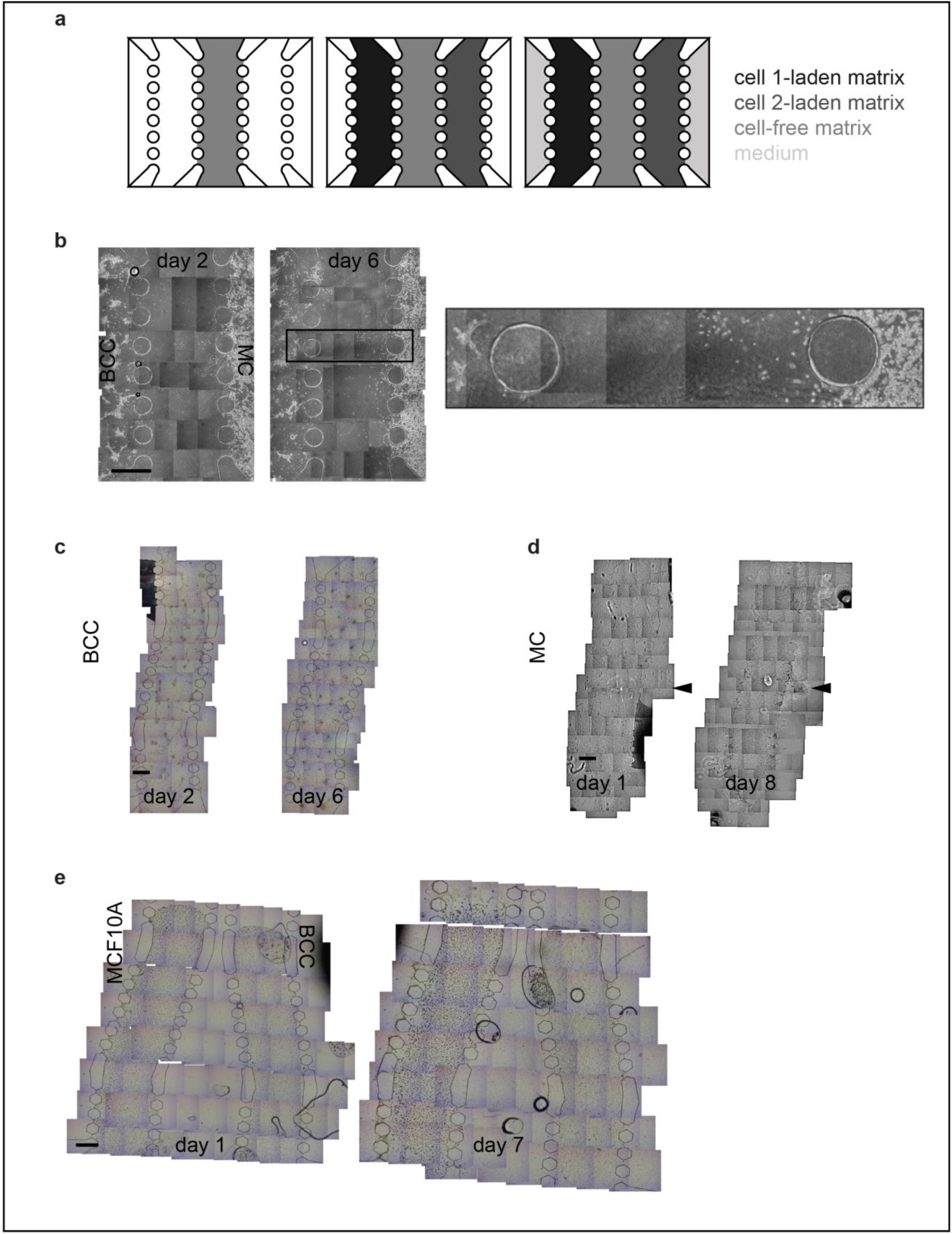
(a) Cell-on-a-chip design to test distant interactions (not drawn to scale). Cell-free matrix was loaded into the middle channel. Cell-laden matrices were loaded into channels on either side of the middle channel. The two reservoirs neighbouring the cell-laden channels were filled with cell culture medium. (b) Representative image for a cell-on-a-chip device where the cell-free middle channel had a constant width (n = 2 cell-on-a-chip devices). Image on the right shows the zoom-in region marked with a rectangle on the image on the left. Representative images of control experiments where only BCC (c) or only macrophages (MC) (d) or BCC across normal mammary epithelial cells (MCF10A) (e) were cultured in the DDI-chip (n = 10 cell-on-a-chip devices). Cell-free matrix was loaded into the middle channel. Cell-laden matrices were loaded into side channels adjacent the middle channel. For only BCC (c) and only MC (d), one side channel was loaded with cell-laden matrix while the other side channel was loaded with cell-free matrix. The leakage of MC which was apparent on day 1 in (d) was marked with black arrowheads. These cells therefore did not really migrate by day 8. In the control chip with BCC and MCF10A (e), cell-laden matrices were loaded into sides channel in the same chip, testing their distant interaction. The two reservoirs neighbouring the cell-laden channels were filled with cell culture medium. Scale bars 1 mm.

## Supplementary Videos

**Supplementary Movie S1**. Simulation of diffusion of fluorescent dextran into the middle channel of the DDI-chip

**Supplementary Movie S2**. EGFR endocytosis in BCC transfected with EGFR-GFP and starved.

**Supplementary Movie S3**. EGFR endocytosis in BCC transfected with EGFR-GFP, starved and treated with EGF.

**Supplementary Movie S4**. EGFR endocytosis in BCC transfected with EGFR-GFP, starved and treated with fluorescently labelled macrophages.

## Supplementary Datasets

**Supplementary Excel File 1**. All statistical test results.

**Supplementary Excel File 2**. All data used for statistical analysis.

